# Cre-dependent optogenetic transgenic mice without early age-related hearing loss

**DOI:** 10.1101/416164

**Authors:** Daniel Lyngholm, Shuzo Sakata

## Abstract

With the advent of recent genetic technologies for mice, it is now feasible to investigate the circuit mechanisms of brain functions in an unprecedented manner. Although transgenic mice are commonly used on C57BL/6J (C57) background, hearing research has typically relied on different genetic backgrounds, such as CBA/Ca or CBA due to the genetic defect of C57 mice for early age-related hearing loss. This limits the utilization of available genetic resources for hearing research. Here we report congenic (>F10) Cre-dependent channelrhodopsin2 (ChR2) mice on CBA/Ca background. By crossing this line with Cre-driver mice on C57 background, F1 hybrids restored the hearing deficit of C57 mice. We also found a linear relationship between aging and hearing loss, with progression rates varied depending on genetic backgrounds (3.39 dB/month for C57; 0.82 dB/month for F1 hybrid). We further demonstrate that this approach allows to express ChR2 in a specific type of inhibitory neurons in the auditory cortex and that they can be identified within a simultaneously recorded population of neurons in awake mice. Thus, our Cre-dependent optogenetic transgenic mice on CBA/Ca background are a valuable tool to investigate the circuit mechanisms of hearing across lifespan.

## Introduction

Recent developments in various genetic tools and technologies have revolutionized the investigation of the circuit level mechanisms underlying various behaviors (Yizhar et al., 2011;Deisseroth and Schnitzer, 2013;Wietek et al., 2014;Buzsaki et al., 2015;Rajasethupathy et al., 2016;Roth, 2016;Blackwell and Geffen, 2017;Jun et al., 2017;Gutruf and Rogers, 2018). While mice increasingly play a crucial role in the advancement of neuroscience research, most research is conducted using the C57BL/6J (C57) mouse strain.

A growing number of hearing researchers have also employed advanced optogenetic technologies developed in C57 mice (Seybold et al., 2015;Nelson and Mooney, 2016;Phillips and Hasenstaub, 2016;Blackwell and Geffen, 2017;Guo et al., 2017;Kato et al., 2017). However, because C57 mice are known to develop early hearing loss from the age of 6-8 months due to a point mutation of *cdh23* gene (Zheng et al., 1999; Noben-Trauth et al., 2003), this poses limitations on hearing research especially when investigating the aging auditory system.

Comparing auditory functions between C57 mice and other genetic backgrounds, such as, CBA or CBA/Ca (CBA) mice, has been a popular approach to study the aging auditory system. This approach allows for dissociation of peripheral and central effects of aging on auditory processing (Frisina, 2001;Frisina et al., 2011). However, because of limited availability of transgenic CBA mice, transgenetic approaches are not straightforward. For example, genetically targeting a specific cell-type in both C57 and CBA backgrounds by utilizing available Cre-driver mice is not currently feasible.

Since C57 × CBA F1 hybrid mice restore the *cdh23* mutation (Frisina et al., 2011), generating transgenic mice on a C57 background and then breeding them with CBA wild-type mice can create a valuable transgenic tool for examining the auditory system without early onset hearing loss. However, if the gene-of-interest is located on the same chromosome as the *cdh23* gene (i.e., chromosome 10), this approach will require additional considerations (such as genotyping for multiple genes) for an appropriate experimental design.

To address this limitation and to broaden the resource for hearing research, here we present Cre-dependent optogenetic transgenic mice on CBA/Ca background (Ai32^cba/ca^) to express channelrhodopsin2 (ChR2) in a cell-type-specific manner. We developed a congenic (>F10) line of Ai32^cba/ca^ mice. By crossing a Cre-driver line on C57 background with the Ai32^cba/ca^ mice (**Figure 1A**), we confirm that 1) ChR2 can be expressed in the auditory cortex of F1 hybrids in a cell-type-specific manner, 2) the F1 hybrids retain hearing threshold at >1 year old compared to transgenic mice on C57 background alone, and 3) ChR2-positive neurons can be identified *in vivo*. Thus, this Ai32^cba/ca^ mouse line allows auditory researchers to utilize a variety of Cre-driver mice to express ChR2 in the auditory system and to facilitate studies of the mouse auditory system, in particular in the context of aging.

**Figure 1:**
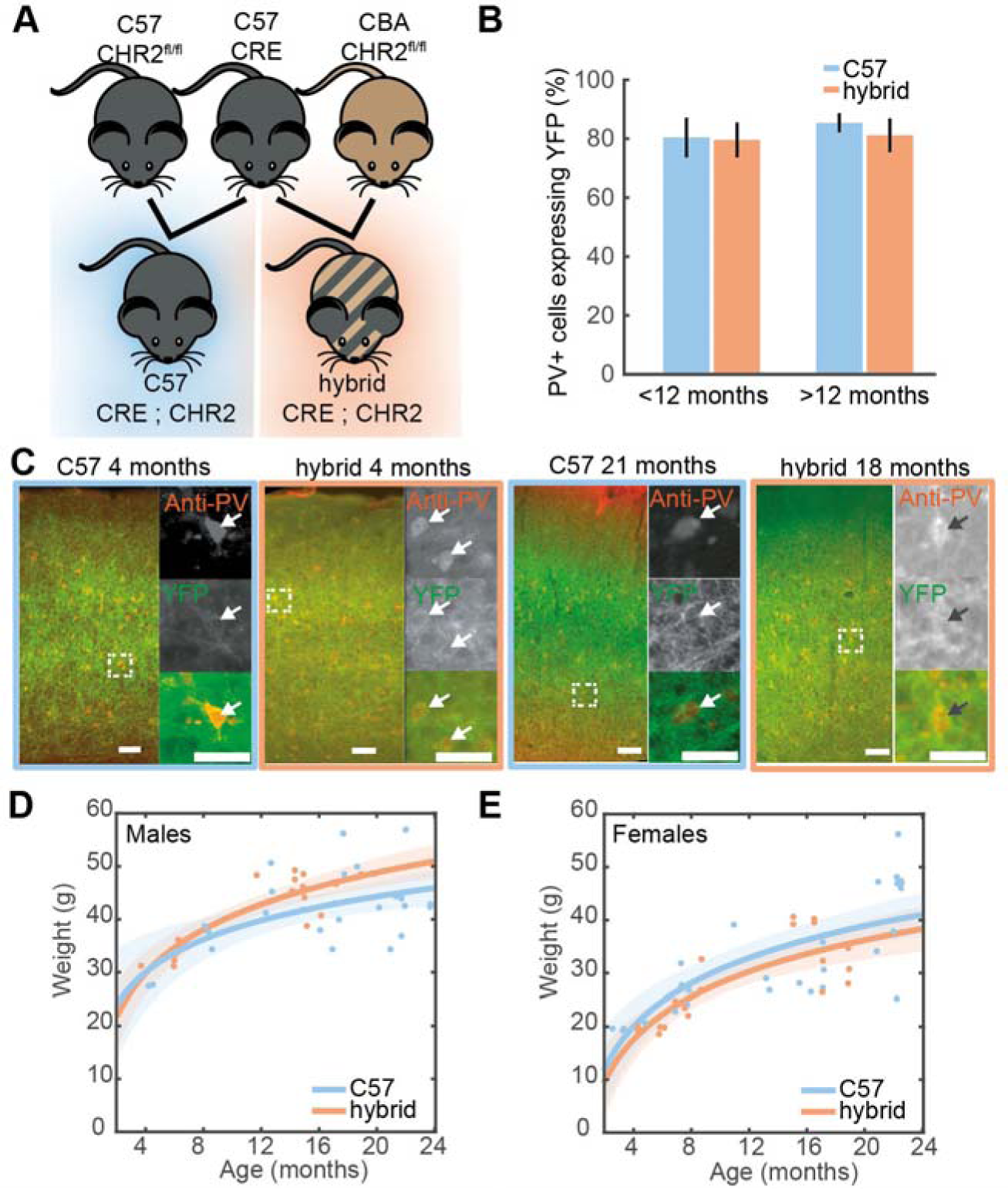
Cell-type-specific expression of ChR2 in the auditory cortex. **A)** A genetic strategy to compensate for the hearing deficit in C57 mice. A Cre-driver line on C57 background (*top middle*) was crossed with Ai32^cba/ca^ mice, a floxed ChR2 mouse line on CBA/Ca background (*top right*), to create F1 hybrid mice (*bottom right*). **B)** Proportion of cells expressing PV that co-expressed ChR2-YFP in the auditory cortex of young (<7 month old) and old (>15 month old) PV-Cre^c57^∷Ai32^c57^(C57) (n = 10) and PV-Cre^c57^∷Ai32^cba/ca^(hybrid) (n = 10) mice (for strain, F(1,19) = 0.28, *p* = 0.61; for age, F(3,19) = 1.25, *p* = 0.35, two-way ANOVA). Error bars represent SEM. **C)** Examples of ChR2-YFP expression (green) stained with anti-PV (red) in the auditory cortex from C57 and hybrid from young (left) and old (right) animals. Enlarged images represent co-expression of PV and ChR2-YFP. Scale bar, 50 µm. **D-E)** Weight at time of recording for male **(D)** (33 C57, 19 hybrids; F(1,44) = 0.05, *p* = 0.92, one-way ANCOVA) and female mice (**E**) (20 C57, 17 hybrids; F(1,39) = 0.79, *p* = 0.38, one-way ANCOVA). Line is an exponential fit (R^2^_C57female_ = 0.56; R^2^_hybrid female_ = 0.64). Shaded area represents the 95% confidence interval.

## Materials and Methods

### Animals

All animal experiments were performed in accordance with the UK Animals (Scientific Procedures) Act of 1986 Home Office regulations and approved by the Home Office (PPL 70/8883). Ai32 mice (JAX012569) (Madisen et al., 2012) have been backcrossed onto either a CBA/Ca background (Ai32^cba/ca^, F18 to date) or a C57Bl/6J background (Ai32^c57^, F10 to date) in house. Mice expressing Cre in either Parvalbumin (PV-Cre) (JAX008069) or Somatostatin (SOM-Cre) (JAX013044) cells were maintained on C57Bl/6J background (≥ F6) and crossed with either Ai32^cba/ca^(≥ F10) or Ai32^c57^(≥ F6) (**Figure 1A**). Genotyping for the genetic background of Ai32 mice was designed to determine whether the nucleotide 753 of *cdh23* gene is adenine or not. All genotyping was performed by Transnetyx using real-time PCR. Ai32^cba/ca^mice tested were all negative whereas Ai32^c57^tested were all positive.

Mice were kept for up to 2 years within the local animal facility. To maintain their body weight, low-calorie diet was given from 3 months of age. In the present study, a total of 89 mice (35 PV-Cre^c57^∷Ai32^c57^; 21 PV-Cre^c57^∷Ai32^cba/ca^; 17 SOM-Cre^c57^∷Ai32^c57^; 16 SOM-Cre^c57^∷Ai32^cba/ca^) were used. Their age and gender for histological (**Figure 1**) and electrophysiological studies (**Figure 2**) are summarized in **Tables 1 and 2**.

**Table 1:** Number of animals used for histological analysis

|  | <12 |  | >12 |  |
| --- | --- | --- | --- | --- |
|  | C57 | F1 hybrid | C57 | F1 hybrid |
| **Female** | 3 | 2 | 2 | 2 |
| **Male** | 2 | 3 | 3 | 3 |

**Table 2:** Number of animals used for establishing hearing threshold

|  | <12 months |  | >12 months |  |
| --- | --- | --- | --- | --- |
|  | C57 | F1 hybrid | C57 | F1 hybrid |
| **Female** | 9 | 9 | 11 | 8 |
| **Male** | 5 | 7 | 28 | 12 |

**Figure 2:**
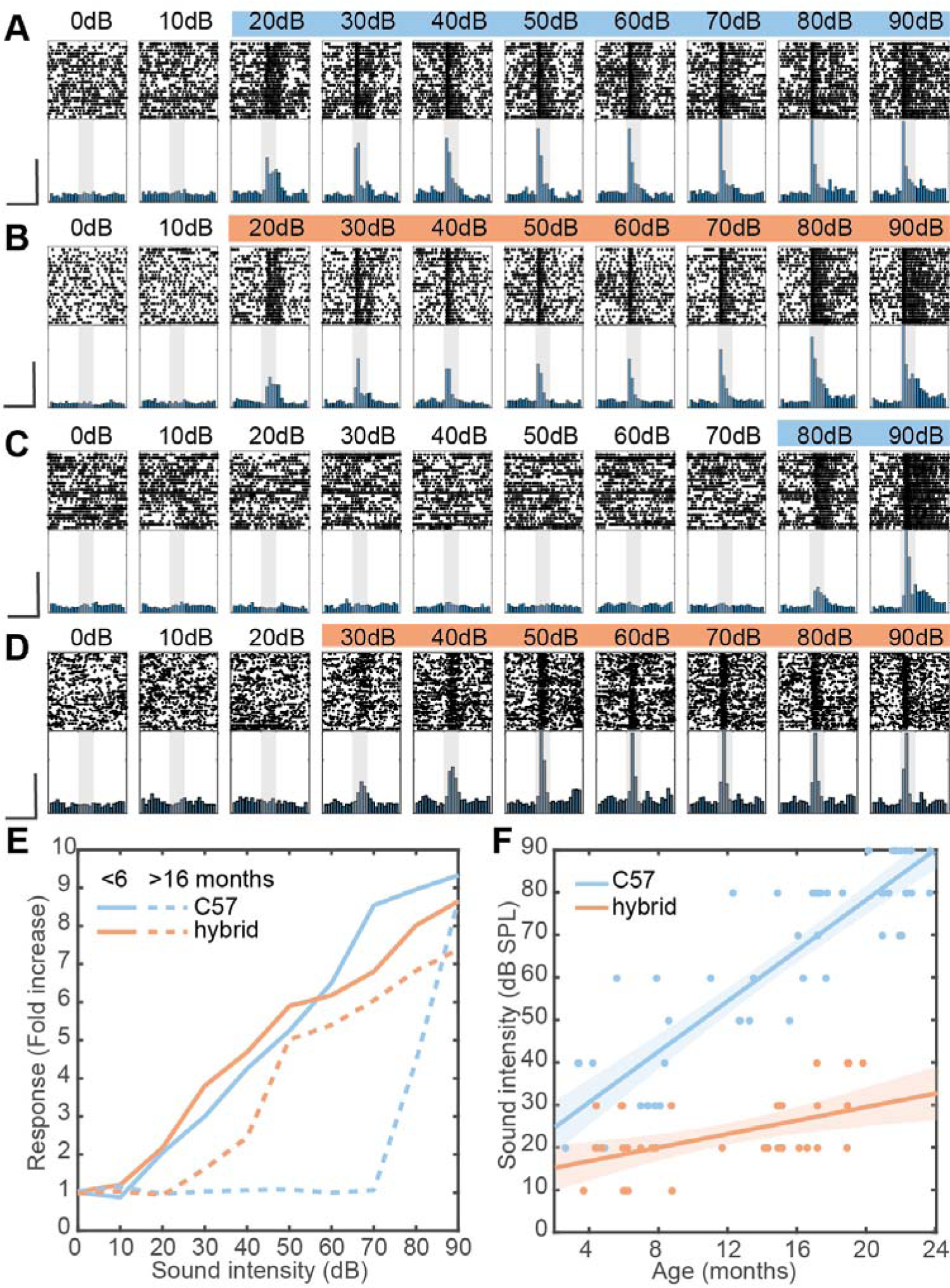
Restoration of hearing deficit in F1 hybrid and linear relationship between aging and auditory threshold. **A-D)** Examples of auditory evoked responses with varied sound intensities in the AC of young C57 (**A**), young hybrid (**B**), old C57 (**C**) and old hybrid mice (**D**). Spike raster (top) and normalized peri-stimulus time histogram (bottom) for MUA are shown. Colored sound intensities elicited statistically significant increase in firing rate over baseline. Shaded area, 100 ms broadband white noise stimulation. Scale bars, 200 ms and 50% of maximum firing rate. **E)** Quantification of exemplar auditory-evoked responses (**A-D**), with the fold change from baseline in firing rate during sound presentation. **F)** Changes in sound intensity threshold as a function of age in C57 (n = 53) and F1 hybrid (n = 38). Data was fitted by linear polynomial functions (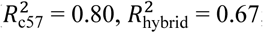; slope_c57_ = 3.39 dB/month, slope _hybrid_ = 0.82 dB/month; T _slope_ = 6.09, *p* _slope_ = 1.46 × 10^-7^, ANCOVA; *y*-intercept_c57_ = 18.37 ± 2.89 dB, *y*-intercept _hybrid_ = 12.38 ± 2.89 dB, T _intercept_ = 1.04, *p* _intercept_ = 0.30, ANCOVA). Shaded area represents the 95% confidence interval.

### Surgery

All procedures have been previously described (Yague et al., 2017). Briefly, animals were anesthetized with isoflurane (1 - 1.5%). Lidocaine (2%, 0.1–0.3 mg) was administered subcutaneously at the site of incision and Carprofen (Rimadyl, 5 mg/kg) was administered intraperitoneally to provide analgesia after the surgery. A head-post was attached on the skull by implanting two frontal bone screws (AP +3 mm, ML 2 mm from bregma), one of which was used for cortical electroencephalogram (EEG) recording. Another two screws were implanted over the cerebellum, one of them used as a ground and a reference. A pair of nuts was then attached with dental cement as a head-post. After the head-post surgery the animals were left to recover for at least 5 days. During an acclimation period of 5 days, the animals were placed in a head-fixed apparatus (SR-8N-S, Narishige), by holding them securely by the head-post and placing the animal’s body in an acrylic tube. This procedure was continued for at least 5 days, during which the duration of head-fixation was gradually extended from 15 to 60 min. During this period, the animals were also exposed to the sound stimulation in the same manner as the electrophysiological recording (see below). A day after this acclimation period, the animals were anesthetized with isoflurane and a craniotomy (2×2 mm^2^at 2.3 mm posterior and 4.2 mm lateral to bregma) was performed to expose the primary auditory cortex (AC). The cranial window was protected with a biocompatible sealant (Kwik-Sil, World Precision Instruments). The following day, the animals were placed in the head-fixed condition for electrophysiological recording.

### *In vivo* electrophysiology

Detailed recording procedures are the same as those described in previous works (McAlinden et al., 2015;Scharf et al., 2016;Yague et al., 2017). All electrophysiological recordings were performed in a single-walled acoustic chamber lined with 3 inches of acoustic absorption foam (MAC-3, IAC Acoustics). Mice were head-fixed and either a 32 or 64 channel silicon probe (A1×32–10 mm–25s–177-A32 or A4×16-10 mm-50s-177-A64, respectively, NeuroNexus Technologies) was inserted using a manual micromanipulator (SM-25A, Narishige) for AC recordings. Probes were inserted at a 40-50° angle to be perpendicular to the cortical surface (800 – 1000 μm depth from the cortical surface). The location of the electrode in AC was assessed by evaluating the local field potential (LFP) and multiunit activities (MUA) in response to white noise stimulation (see below).

Broadband signals were amplified (RHD2132, Intan Technologies, LLF) relative to the ground and were digitized at 20 kHz (RHD2132 and RHD2000, Intan Technologies, LLC). The recording session was initiated >30 min after the probe was inserted to its target depth, to allow for signal stabilization. A typical recording session consisted of >15 min baseline recording of spontaneous activity, followed by an optical stimulation protocol, sound presentation and then another baseline of spontaneous activity.

### Optical stimulation

Pulses of blue light (450 nm, PlexBright, Plexon) of 100 ms duration were delivered at 2 Hz through a 200 µm fiber optic (Plexon) attached to the silicon probe and positioned on the surface of the brain. The light output at tip of the fiber optic was measured with a constant long (> 1 sec) light pulse before probe insertion and was 45 ± 14 mW/mm^2^(mean ± SD).

### Sound presentation

Sound was generated digitally (sampling rate 97.7 kHz, TDT, Tucker-Davis Technologies) and delivered in free-field through a calibrated electrostatic loud-speaker (ES1) located ∼15 cm in front of the animal. To estimate the hearing threshold of animals, broadband white noises (100 ms with 5 ms cosine ramps, 10 dB steps, 0-90 dB SPL) were pseudo-randomly presented with a minimum of 400 ms interval for 25 repetitions.

### Histology

For verification of silicon probe tracks, the rear of probes was painted with DiI (∼10% in ethanol, D282, Life Science Technologies) before probe insertion. After electrophysiological experiments, animals were perfused transcardially with physiological saline followed by 4% paraformaldehyde/0.1 M phosphate buffer, pH 7.4. After an overnight post-fixation in the same fixative, brains were stored in 30% sucrose in phosphate buffered saline (PBS) for cryoprotection. Brains were then cut into 50 μm coronal sections with a sliding microtome (SM2010R, Leica) and placed in PBS.

To visualize parvalbumin-positive (PV+) neurons in PV-Cre^c57^∷Ai32^c57^and PV-Cre^c57^∷Ai32^cba/ca^, immunohistochemistry was also performed. After slicing, a subset of sections were incubated with a blocking solution (10% normal goat serum, NGS, in 0.5% Triton X in PBS, PBST) for 1 hour at room temperature followed by incubating primary antibodies (anti-PV 1:4000, P3088, Sigma-Aldrich) in 3% NGS in PBST at 4 °C overnight. After washing, sections were incubated with secondary antibodies (Goat anti-mouse Alexa Fluor 568, 1:500, A11007, Life Science Technologies) for 2 hours at room temperature. After washing, sections were mounted on gelatin-coated slides and cover-slipped with antifade solution. Sections were also stained with DAPI (1µg/ml; Sigma-Aldrich) to determine cortical laminae and structural landmarks used to aid localization of the AC. The sections were mounted on gelatin-coated slides and cover-slipped with antifade solution (Vectashield, Vector Laboratories).

### Data analysis

Data analysis was performed offline using MATLAB (Mathworks) or freely available software. To extract local field potentials (LFPs), a lowpass filter (<100 Hz) was applied and signals were downsampled to 1 kHz. For spike detection and sorting, the Klusta package (Rossant et al., 2016) or Kilosort (Pachitariu et al., 2016) was used. During visual inspection after this automatic process, events that occurred across all channels were excluded as noise. Other clusters were categorized as either single-unit or multi-unit activity. The quality of clusters was further assessed by measuring isolation distance (Schmitzer-Torbert et al., 2005). The inclusion criteria for single units were ≥30 isolation distance and ≥0.1 Hz spontaneous firing. Below, multi-unit activity (MUA) includes both single-unit and multi-unit clusters.

To estimate the hearing threshold of mice, the MUA firing rates during a 50 ms before and after onset of stimulation were compared. Significance was determined using a Bonferroni corrected Wilcoxon signed-rank test with a 5% significance level threshold. The hearing threshold was designated as the lowest sound intensity which resulted in a significant increase in the median firing rate. Differences in the hearing threshold between two genetic backgrounds were assessed using two-way analysis of covariance (ANCOVA), with respect to either slope or *y*-intercept.

To determine whether single units were narrow or broad spiking cells, the trough-to-peak duration and width at 20% of spike amplitude of averaged spike waveforms were computed. Since these measures showed a bimodal distribution, single units were classified into 2 clusters using spectral k-means clustering implemented using spectral clustering from the Python scikit-learn library (Pedregosa et al., 2011).

To identify single units modulated by optogenetic stimulation in PV-Cre^c57^∷Ai32^c57^and PV-Cre^c57^∷Ai32^cba/ca^mice (**Figure 3**), firing rates were compared during a 20 ms window before and after stimulation using a Bonferroni corrected Wilcoxon signed-rank test with a 5% significance level threshold. Units with a significant increase in the median firing rate during stimulation were classified as positively modulated (presumptive PV+ cells) and units with a significant decrease were classed as negatively modulated cells.

**Figure 3:**
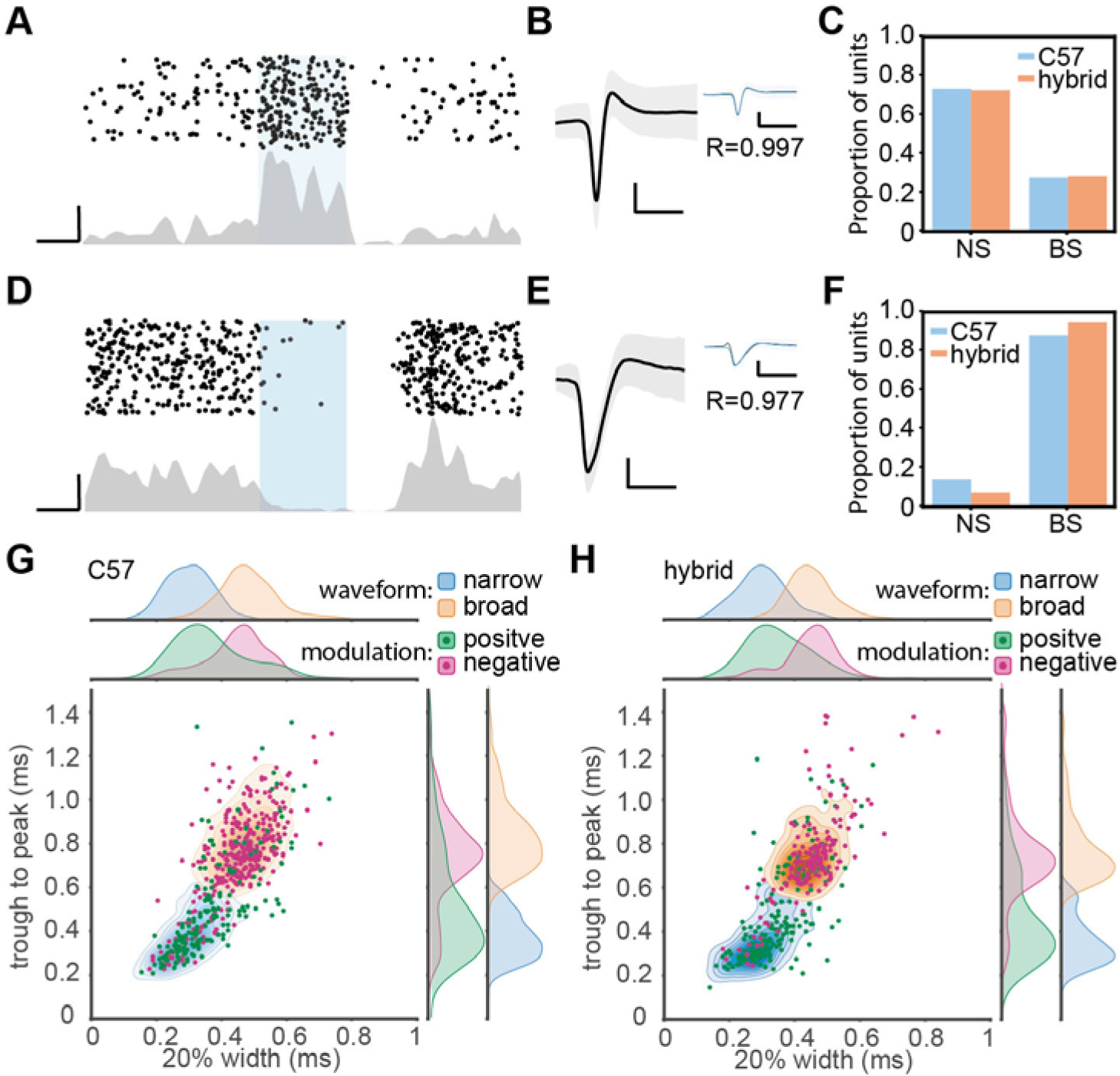
Optogenetic manipulation of auditory cortical PV+ neurons. **A)** Raster plot (*top*) and peri-stimulus time histogram (*bottom*) for a single unit showing significant positive modulation by optical stimulation in a PV-Cre^c57^∷Ai32^cba/ca^mouse. Scale bars, 50 ms and 20 Hz. **B)** Average waveform for the same single unit shown in **A**. Shaded area represents the standard deviation. Inset shows the overlay of average waveform with (blue) and without (black) optical stimulation. *R* represents the correlation coefficient between the two average waveforms. Scale bars, 1 ms and 0.1 mV. **C)** The proportion of narrow spiking (NS) and broad spiking (BS) cells within optically activated cells in C57 (blue) and hybrid (orange) animals. **D-F)** Same as A-C, but for optically suppressed cells. **G and H)** Scatter and kernel density plot of trough-to-peak during and 20% width of spike waveforms across single units in C57 (996 single units from 36 PV-Cre^c57^∷Ai32^c57^) (**G**) and hybrid (759 single units from 20 PV-Cre^c57^∷Ai32^cba/ca^) (**H**). Distributions of NS and BS cells were shown as a density plot (light blue, NS cells; light orange, BS cells), with each density level representing 10% of the population. *Green*, optically activated cells. *Pink*, optically suppressed cells. Distribution of each spike waveform measurement for each cell class was also estimated by using a kernel function analysis.

The density functions in **Figures 3G and H** were estimated by fitting a Gaussian kernel to the data as implemented in the Python scikit-learn library (Pedregosa et al., 2011).

### Statistical analysis

Data was presented as mean ± SEM unless otherwise stated. Statistical analyses were performed with the Python Statsmodels package. For fitted regression lines, the shaded area around line represents the root mean squared error of the fitted function. In **Figure 1B**, a two-way ANOVA was performed for age and strain using post-hoc Tukey’s Honestly Significant Difference (HSD) test. In **Figures 1D & E**, a logarithmic fit was performed. To test if regression was different between strains, an ANCOVA with post-hoc HSD test was performed on log transformed data. In **Figure 2F**, a linear fit was performed. To compare regression lines, an ANCOVA with post-hoc HSD test was performed. **Figures 3G & H**, a one-way ANOVA was performed and effect size was reported as eta-squared (η^2^).

## Results

### Database

To determine the threshold of neural responses to white noise with varied intensities (**Figure 2**), we analyzed 111 recordings from 89 animals (**Table 2**). For *in vivo* opto-electrophysiological experiments (**Figure 3**), we isolated and analyzed single units from a subset of the above, comprising 53 recordings from 36 PV-Cre^c57^∷Ai32^c57^ mice and 24 recordings from 20 PV-Cre^c57^∷Ai32^cba/ca^ mice (**Table 3**). A further subset of 20 PV-Cre^c57^∷Ai32^c57^ mice (**Table** 1) were used to analyze the expression of PV (**Figure 1**).

**Table 3:** Number of single units used for waveform analysis

|  | Positive modulation |  | Negative modulation |  | No modulation |  |
| --- | --- | --- | --- | --- | --- | --- |
|  | C57 | F1 hybrid | C57 | F1 hybrid | C57 | F1 hybrid |
| **BS** | 50 | 51 | 150 | 79 | 419 | 335 |
| **NS** | 133 | 131 | 24 | 6 | 220 | 157 |

### Cell-type-specific expression of ChR2 in the auditory cortex

We examined expression of ChR2-EYFP in the auditory cortex (AC) on both genetic backgrounds at different ages (**Figure 1C**). In young PV-Cre^c57^∷Ai32^c57^ mice (3-7 month old), we confirmed that 80.4 ± 6.7% of PV+ neurons expressed ChR2-EYFP in the AC. This trend was held with 85.4 ± 3.3% in older animals (15-24 month old) (**Figure 1B**). Similarly for PV-Cre^c57^∷Ai32^cba/ca^ mice, 79.6 ± 5.9% and 81.2 ± 5.7% of PV+ neurons expressed ChR2-EYFP in the AC for young and old mice, respectively. We observed no significant differences of expression in either strain (F(1,19) = 0.28, *p* = 0.61, two-way ANOVA) or age (F(3,19) = 1.25, *p* = 0.35, two-way ANOVA) (**Figure 1B**). We observed no differences in the weight gain between both backgrounds in either female (R^2^_C57_ female = 0.56; R^2^_hybrid female_ = 0.64) (F(1,44) = 0.05, *p* = 0.92, one-way ANCOVA) or male mice (R^2^C57 male = 0.75; R^2^hybrid male = 0.68) (F(1,39) = 0.79, *p* = 0.38, one-way ANCOVA) (**Figures 1D, E**). Thus, both genetic mice are comparable with respect to the weight gain across age and more importantly, Ai32^cba/ca^ mice allow expression of ChR2 in a Cre-dependent manner in the AC across age.

#### Linear relationship between aging and auditory threshold with varied rates

Early age-related hearing loss in C57 mice is due to a mutation in the *cdh23* gene (Noben-Trauth et al., 2003) and a previous report showed crossing C57 mice with mice on CBA background can restore the hearing loss (Frisina et al., 2011). To confirm whether the F1 hybrid between Cre-driver mice on C57 and Ai32^cba/ca^ can restore the hearing deficit, we compared auditory cortical evoked responses to broadband white noise with varied sound intensities between both C57 background and the F1 hybrid across age under a head-fixed unanesthetized condition (**Figure 2**). We took multi-unit activity (MUA) to estimate the sound intensity threshold.

Representative examples of sound-evoked responses are shown in **Figures 2A-E**. The response profiles across intensities were comparable between the two young mice (3-4 months old) on the two backgrounds (**Figures 2A, B**). However, a 15 month old C57 mouse showed the detrimental effect of aging on auditory evoked responses (**Figure 2C**), while a 16 month old F1 hybrid still showed robust evoked responses at lower intensities (**Figure 2D**). To confirm these trends, we quantified auditory evoked responses across intensities for each animal (**Figures 2E**).

To further investigate this trend across animals (**Table 2**), we assessed the sound intensity threshold as a function of age (**Figure 2F**). Although the initial hearing threshold at 3-4 month old was similar between both strains, threshold progressively increased in C57 mice over 2 years. In contrast, F1 hybrid mice showed smaller changes over the same period (**Figure 2F**). To quantitatively assess these trends of age-related hearing loss, we fitted data by a simple linear model. While the *y*-intercept for C57 (18.37 ± 2.89 dB) and F1 hybrid (12.38 ± 2.89 dB) was not significantly different between genetic backgrounds (T = 1.04, *p* = 0.30, ANCOVA), the slope of C57 mice was significantly higher than that of F1 hybrids (3.39 dB/month for C57 vs 0.82 dB/month for F1 hybrids) (T = 6.09, *p* = 1.46 × 10^-7^, ANCOVA). Therefore, crossing Cre-driver mice on a C57 background with Ai32^cba/ca^ can significantly diminish the effect of the genetic defect.

### Optogenetic manipulation of PV+ neurons *in vivo*

To utilize Ai32^cba/ca^ mice for *in vivo* optogenetic experiments, we applied optical stimulation to the AC of head-fixed awake PV-Cre^c57^∷Ai32^c57^ (C57, n = 36) and PV-Cre^c57^∷Ai32^cba/ca^ mice (F1 hybrid, n = 20) while auditory cortical ensembles were electrophysiologically monitored by inserting a silicon probe.

Of 1755 single units (996 cells from C57; 759 cells from F1 hybrids), we classified cells based on optical evoked responses (**Figures 3A-F**) and spike waveforms (**Figures 3G, H**). **Figures 3A** and **3D** show two examples of optically activated and suppressed cells, respectively, together with their spike waveforms (**Figures 3B, E**). In total, we identified 365 (20.8 %) optically activated cells (183 cells from C57; 182 cells from F1 hybrid) and 259 (14.8 %) optically suppressed cells (174 cells from C57; 85 cells from F1 hybrid). The former cells are presumably PV+ neurons.

In parallel, we also classified cells into broad spiking (BS) and narrow spiking (NS) cells based on spike waveforms by measuring trough-to-peak duration (T2P) and the 20% width of the spike deflection (W20) (**Figures 3G, H and Table 3**). We obtained 1152 (65.6 %) BS cells (653 cells from C57; 499 cells from F1 hybrid) and 603 (34.4 %) NS cells (343 cells from C57; 260 cells from F1 hybrid).

Based on these two classification approaches, we assessed the proportion of BS and NS cells within optically activated (**Figure 3C**) and suppressed cells (**Figure 3F**). In C57 mice, 65.0 % (119/183) and 35.0 % (64/183) of optically activated cells are NS and BS cells, respectively (**Figure 3C**). F1 hybrid mice also showed a similar trend (123/182 for NS cells; 59/182 for BS cells). With respect to optically suppressed cells, the majority of them were BS cells (157/174 in C57; 81/85 for F1 hybrid) (**Figure 3F**). The mean of spike waveform features was similar in both strains for both T2P (F(1,1751) = 0.44, *p* = 0.75, η^2^= 0.0002, one-way ANOVA) and W20 (F (1,1751) = 3.37, *p* = 0.13, η^2^= 0.002, one-way ANOVA), indicating that spike waveform features are comparable between strains. Thus, Ai32^cba/ca^ mice can be also used for *in vivo* optogenetic experiments.

## Discussion

To utilize recent genetic technologies in mice for hearing research, we presented Cre-dependent ChR2 mice on CBA/Ca background, Ai32^cba/ca^ mice. By crossing this line with Cre driver mice on C57 background, we demonstrated 1) cell type-specific expression of ChR2 in the AC, 2) the restoration of early age-related hearing loss in C57 mice, and 3) the capability of optogenetic manipulations *in vivo*. Thus, this transgenic line offers an opportunity to investigate age-related changes in auditory functions in a cell type-specific manner.

In the present study, we assessed the threshold of auditory evoked responses based on MUAs in the AC. Our results generally agree with the previous report, which examined the same F1 hybrid using auditory brainstem response (ABR) (Frisina et al., 2011) and are broadly in line with previous observations using ABR in CBA/Ca background (Kane et al., 2012). In addition to replicating the previous finding, we also found a clear linear relationship between age and hearing threshold for the first time (**Figure 2E**). This linear relationship suggests that age-related hearing loss in C57 mice is an incremental process rather than a sudden change to pathophysiological conditions at a certain time point. Because the effect of the genetic defect seems to be apparent even in young (< 12 month old) C57 mice, we recommend that studies always report the genetic background. It is important to replicate our results with a larger sample size as well as different assessments including ABR and tone evoked responses across frequencies in the future.

While we have developed Cre-dependent optogenetic mice on CBA/Ca background, there are other approaches to restore the genetic defect of C57 mice. One complementary approach is to generate transgenic mice expressing the gene of interest on the C57 background, and then breed them with mice without the point mutation of *cdh23* gene. In addition to wild-type CBA or CBA/Ca mice, B6.CAST-Cdh23^Ahl+^ mice can be used because they possess a wild-type *cdh23* locus (Keithley et al., 2004). Alternatively, mice created by gene editing technology can be an option. In B6-Cdh23^c.^753^G^ mice, for example, the point mutation of *cdh23* has been corrected (Johnson et al., 2017).

One caveat for this alternative approach is that if the gene-of-interest is located on the same chromosome as the *cdh23* gene (i.e., chromosome 10), this approach will require an extra assessment of their genotype. An example of this is vasoactive intestinal peptide gene (VIP). VIP-positive neurons are well known as a prominent subtype of GABAergic interneurons in the cortex (Tremblay et al., 2016). Therefore, to express ChR2 in VIP-positive neurons, crossing VIP-Cre mice with Ai32^cba/ca^ will be a more practical option.

An obvious application of our Ai32^cba/ca^ mice is for aging research. This genetic line allows to investigate how aging affects the central auditory system at the neural circuit level without early age-related hearing loss. Because C57 and CBA mice have been compared to dissociate between peripheral and central effects of aging processing on the auditory system (Frisina, 2001;Frisina et al., 2011), the same approach can be taken together with optogenetic approaches. Thus, our Ai32^cba/ca^ mouse line offers an additional toolbox to investigate age-related and cell type-specific changes in aging auditory system.

## Author contributions

DL and SS designed and conceived experiments. DL performed all experiments and data analysis. DL and SS wrote the manuscript.

## Acknowledgements

We thank Amisha Patel and Caroline Wilson for their comments on an early version of the manuscript.

